# Development of a shortwave infrared (SWIR) sinuscope for the detection of cerebrospinal fluid (CSF) leaks

**DOI:** 10.1101/2022.12.22.520937

**Authors:** Tjadina-W. Klein, Stella Yang, Mahbuba A. Tusty, Jayakar V. Nayak, Michael T. Chang, Oliver T. Bruns, Thomas S. Bischof, Tulio A. Valdez

**Affiliations:** Helmholtz Pioneer Campus, Helmholtz Zentrum München, Neuherberg, Germany; Medizinische Fakultät, Ludwig-Maximilians-Universität München, Munich, Germany; Department of Otolaryngology, Head and Neck Surgery, Stanford University, Palo Alto, California, USA; Department of Functional Imaging in Surgical Oncology, National Center for Tumor Diseases (NCT/UUC) Dresden, Dresden, Germany; Carl Gustav Carus Faculty of Medicine, TU Dresden, Dresden, Germany; Department of Medicine, Technische Universität München, Munich, Germany

**Keywords:** skull base surgery, shortwave infrared, SWIR, sinuscopy, endoscopy, CSF leak

## Abstract

**Significance:** CSF rhinorrhea (leakage of brain fluid from the nose) can be difficult to identify and currently requires invasive procedures such as intrathecal fluorescein which requires a lumbar drain placement. Fluorescein is also known to have rare but significant side effects including seizures and death. As the number of endonasal skull base cases increase, the number of CSF leaks have also increased for which an alternative diagnostic method would be highly advantageous to patients.

**Aim:** To develop an instrument to identify CSF leaks based on water absorption in the SWIR without the need of intrathecal contrast agents. This device needed to be adapted to the anatomy of the human nasal cavity while maintaining low weight and ergonomic characteristics of current surgical instruments.

**Approach:** Absorption spectra of CSF and artificial CSF were obtained to characterize the absorption peaks that could be targeted with SWIR light. Different illumination systems were tested and refined prior to adapting them into a portable endoscope for testing in 3D printed models and cadavers for feasibility.

**Results:** We identified CSF to have an identical absorption profile as water. In our testing, a narrow band laser source at 1480nm proved superior to using a broad 1450 nm LED. Using a SWIR enabling endoscope set up, we tested the ability to detect artificial CSF in a cadaver model.

**Conclusions:** An endoscopic system based on SWIR narrow band imaging can provide an alternative in the future to invasive methods of CSF leak detection.

## 1 Introduction

The diagnosis of cerebrospinal fluid (CSF) leaks can be challenging, and a missed or delayed diagnosis can lead to potentially life-threatening complications such as meningitis (Essayed et al., 2018) (Brown et al., 2018) (Seth et al., 2010). With recent advancements in endoscopic skull base surgery, the incidence of CSF leaks continues to rise, with as high as 30.1% of cases having an intraoperative CSF leak, making iatrogenic causes the leading etiology (Karnezis et al., 2016). In this setting, and particularly in the acute postoperative period, CSF rhinorrhea can be difficult to distinguish clinically from blood and mucus drainage. Other scenarios where a CSF leak may be difficult to discern and locate are in the presence of craniofacial trauma and spontaneous CSF leaks (Schievink, 2008).

Existing options for CSF leak diagnosis such as intrathecal fluorescein or Beta2 transferrin have limitations in immediacy of results and introduce potential morbidity (Lobo, Baumanis, & Nelson, 2017). One possible method to improve diagnostic ability for CSF leak involves shortwave infrared radiation technology (SWIR). We refer to SWIR as the electromagnetic spectrum ranging from 1000 to 2000 nanometers (nm) which is not visible to the human eye and conventional silicon-based cameras (Thimsen, Sadtler, & Berezin, 2017). Compared to traditional visible and near-infrared (NIR) fluorescence imaging, SWIR is capable of providing greater contrast, sensitivity, and penetration depths. Moreover, tissue components such as water, lipids, and collagen have more prominent SWIR absorption features than corresponding features seen in the visible and near-infrared (Wilson, Nadeau, Jaworski, Tromberg, & Durkin, 2015). Still so, SWIR has not yet been readily adapted to clinical settings or diagnostics. This is partly because suitable detectors, until recently, have been either inaccessible or cost prohibitive. Recent studies have highlighted the potential of SWIR technology in clinical diagnostics (Tversky & MacGlashan, 2020). Fluid in the middle ear, for example, shows strong light absorption between 1,400 and 1,550 nm. As a result, straightforward middle ear fluid detection in a model using a SWIR otoscope is possible and can be used to diagnose otitis media with greater accuracy than is currently possible with a standard otoscope (Carr et al., 2018) (Kashani et al., 2021).

In this article, we describe the development of an endoscopic surgical system for assessment of CSF leaks of the skull base, based on SWIR imaging. To begin, we describe the physical principles behind the contrast mechanism and estimate potential performance using in vitro characterization. We then develop a benchtop imaging system and perform proof-of-principle experiments. After testing the initial endoscope design we identify points which can be optimized. We show that the detection quality can significantly be improved when moving to a narrower illumination band and adding polarizers to the design. Using a porcine nasal cavity, we show that the improved design is capable of detecting very small amounts of CSF in a nasal cavity.

## 2 Methods

### 2.1 System/ Design

We based our SWIR endoscope on the design of a regular rigid white light surgical endoscope. A Hawkeye Pro Slim endoscope (Gradient Lens Corporation) was optimized for the wavelength range 900nm to 1700nm using a custom coating. An off the shelf video coupler (VC-35, Gradient Lens Corporation) was modified by exchanging the original lens for a SWIR compatible lens. The SWIR camera was attached to the video coupler using a c-mount extension tube. Two different SWIR cameras were used for this project: the Goldeye G-032 Cool TEC2 (Allied Vision) and the Owl 640 Mini VIS-SWIR (Raptor Photonics). Two polarizers were incorporated into the endoscope. One was positioned at the tip of the endoscope (#12-473, Edmund Optics), the other was positioned in the video coupler in front of the lens (LPNIR050, Thorlabs). Using a laser cutter the polarizer at the tip of the endoscope was custom cut to fit exactly over the fiber ring of the endoscope without covering the lens of the endoscope. The polarizers were oriented orthogonal to each other to minimize direct reflections. The positioning of the polarizers can be seen in figure 1.

**Fig. 1.**
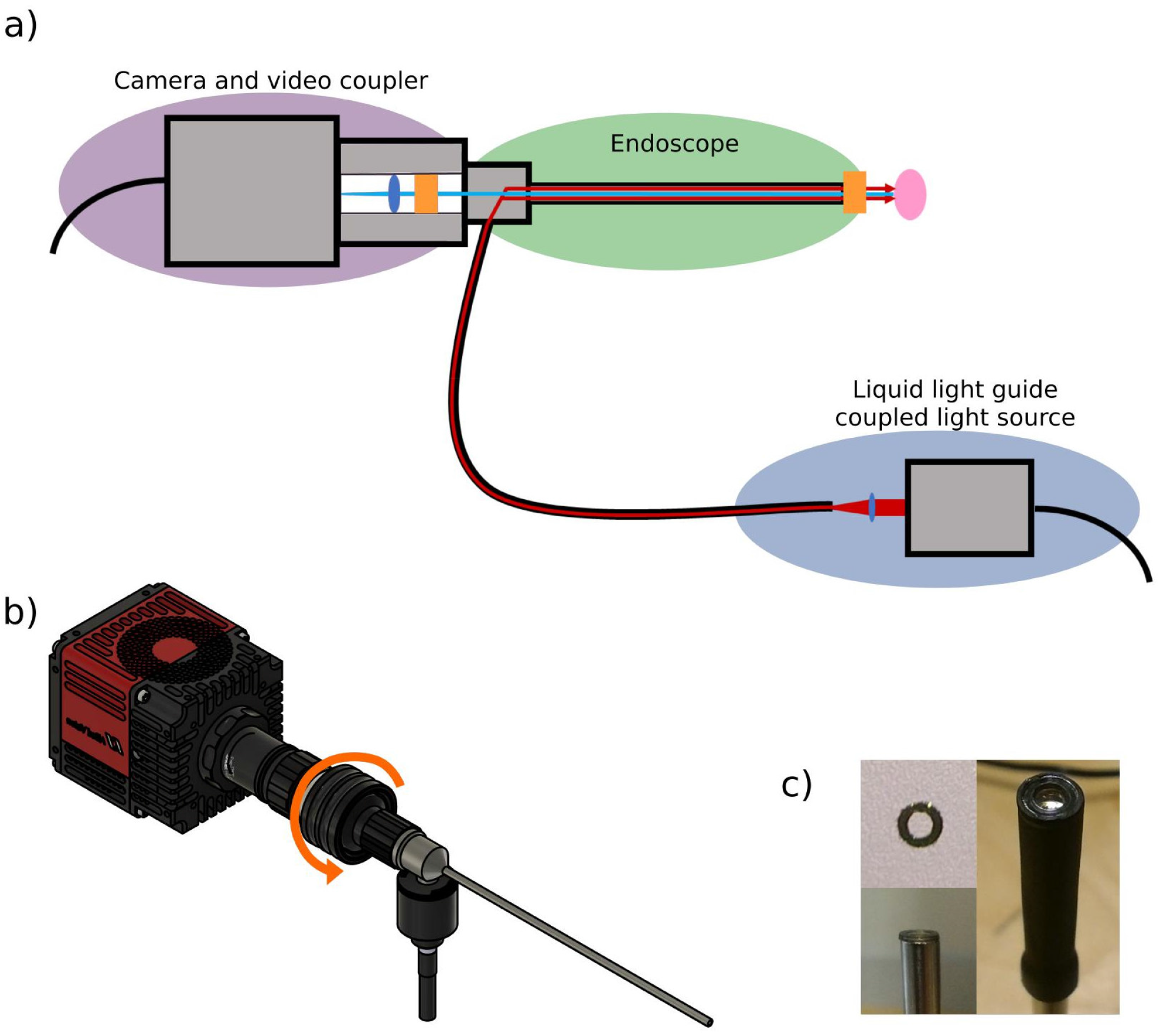
Design of the SWIR endoscope. a) Schematic of the SWIR endoscope, consisting of an InGaAs imager, a video coupler containing a lens and polarizer, a SWIR-optimized rigid endoscope with a second polarizer, and a liquid light guide coupled light source. The blue ellipses and orange rectangles indicate the positions of the lenses and polarizers. The red lines show the path of the illumination beam whereas the blue line represents the light reflected off the sample and collected by the camera. b) CAD design of the SWIR endoscope. The orange arrow shows the part of the setup that can be rotated to align the two polarizers. c) Images of the polarizers used for the endoscope setup: the custom cut polarizer, top view; positioning of the polarizer on the tip of the endoscope; the polarizer covering only the fiber ring and not the lens, held in position by a sleeve (top to bottom, left to right).

The illumination for the endoscope consists of a narrow band 1480nm laser (L1480G1, Thorlabs) which is coupled into the endoscope using a liquid light guide (LLG3-4Z, Thorlabs). A lens (C060TMD-C, Thorlabs) was used to focus the laser into the liquid light guide.

### 2.2 Comparison of white light SWIR performance for visualizing CSF

Transmission spectra were taken of artificial CSF (aCSF) (EcoCyte Bioscience) and of water. aCSF has an electrolyte concentration closely matching CSF with final ion concentrations of NaCl 125mM, KCl 3.0mM, CaCl2 2.5mM, MgSO4 1.3mM, NaH2PO4 1.25mM along with high purity water. 700 microliter volumes of either aCSF or water were transferred to 2mm pathlength quartz cuvettes (Thorlabs CV10Q7F) to measure attenuation spectra using Lambda 1050+ spectrophotometer (Perkin Elmer).

To investigate the potential of the SWIR absorption band seen in the spectroscopic data for fluid detection in imaging data we imaged three Eppendorf tubes, one partially filled with water, one partially filled with aCSF and one empty. They were placed next to each other on a white background and illuminated and imaged from above. The white light image was taken using a white light LED (MNWHL4, Thorlabs), an IDS camera (UI-3880SE-C-HQ) and a Navitar lens (Navitar HR F1.4/8mm). The SWIR image was taken using the Goldeye G-032 Cool TEC2 camera, a Navitar SWIR-35 lens, and a broad band 1450nm LED (M1450L3, Thorlabs).

The SWIR setup used for imaging the water on the chicken skin was identical to that used to image the Eppendorf tubes. For the white light imaging of the chicken skin an IDS camera (UI-3880SE-C-HQ) was used, in combination with a Navitar lens (Navitar HR F1.4/8mm) and a white light LED for illumination (MNWHL4, Thorlabs).

### 2.3 Cadaver study

To evaluate ease of use and proof of principle for visualization of CSF on the skull base, we performed a cadaver study. A trephine was performed through an infrabrow incision to access the frontal sinus and a communication was achieved with the intracranial compartment by puncturing the posterior table of the frontal sinus. A catheter was then inserted into the defect. An intra-nasal injury to the skull base was performed allowing communication for the artificial CSF. The aCSF pumped from the frontal sinus could be assessed from the nasal cavity with our endoscope.

### 2.4 Use of a narrow-band SWIR laser improves CSF contrast

The initial test to assess the effect narrowing the illumination band has on our imaging data was done using the macro setup described above. As a sample we used a petri dish filled with a small amount of water and placed above a test target. To compare relatively broad-band to narrow-band illumination, we performed imaging using a 1450nm LED (M1450L3, Thorlabs) to the same LED with an excitation filter (1480nm center, 12nm width; FB1480-12, Thorlabs). The depth of the water was roughly 1mm (slightly less in the middle of the petri dish).

For the assessment in the endoscopic setup we again used a partially water filled petri dish above a test target. This time the petri dish was slightly angled to create a gradient of water thickness. A very small amount of dish soap was added to the water to reduce the surface tension and create a thinner film of water. The maximum thickness of water in the images is around 1mm, the thinnest part is <<1mm. These images were taken without the use of polarizers. For the broadband illumination, we used a 1450nm LED (M1450L3, Thorlabs), for the narrow band illumination we used a 1480nm laser (L1480G1, Thorlabs).

### 2.5 Reducing reflections through the use of polarizers

One polarizer was positioned inside the video coupler (between endoscope and camera) and a second polarizer was attached to the tip of the endoscope. The polarizer at the tip of the endoscope were custom cut using a laser cutter in order to only cover the fiber ring of the endoscope and not the lens. The final dimensions of the polarizers were 4.3mm (outer diameter) and 2.5mm (inner diameter). The polarizers were oriented orthogonal to each other to bring the reflections to a minimum. Images were taken using the endoscopic setup (first without polarizers, then with polarizers added in) and the sample was a piece of chicken skin. Water was dripped onto the chicken skin during imaging.

### 2.6 Imaging the nasal cavity

To assess our system in a realistic nasal environment we imaged a porcine nose using the improved endoscope system. The endoscope system consisted of the narrow band laser light source and the polarizers, as described above (section 2.1). Our sample was a food grade half pig head. The pig head had been cut in half along the sagittal plane, making it possible to drip water directly into the nasal canal. The endoscope was inserted into the nasal canal and water was dripped into the nasal canal so that it drained towards the endoscope and the tip of the nose.

## 3 Results

### 3.1 Comparison of white light SWIR performance for visualizing CSF

To test our hypothesis that the absorption properties of CSF are similar to water and to predict whether a SWIR endoscope could be beneficial for CSF detection we investigated the optical properties of CSF. The spectroscopic analysis confirmed that the properties of aCSF match those of water for the range of 400nm to 1800nm. The suspected absorption band of aCSF at around 1450nm was clearly visiblein the spectroscopic data (Figure 2a).

**Fig. 2.**
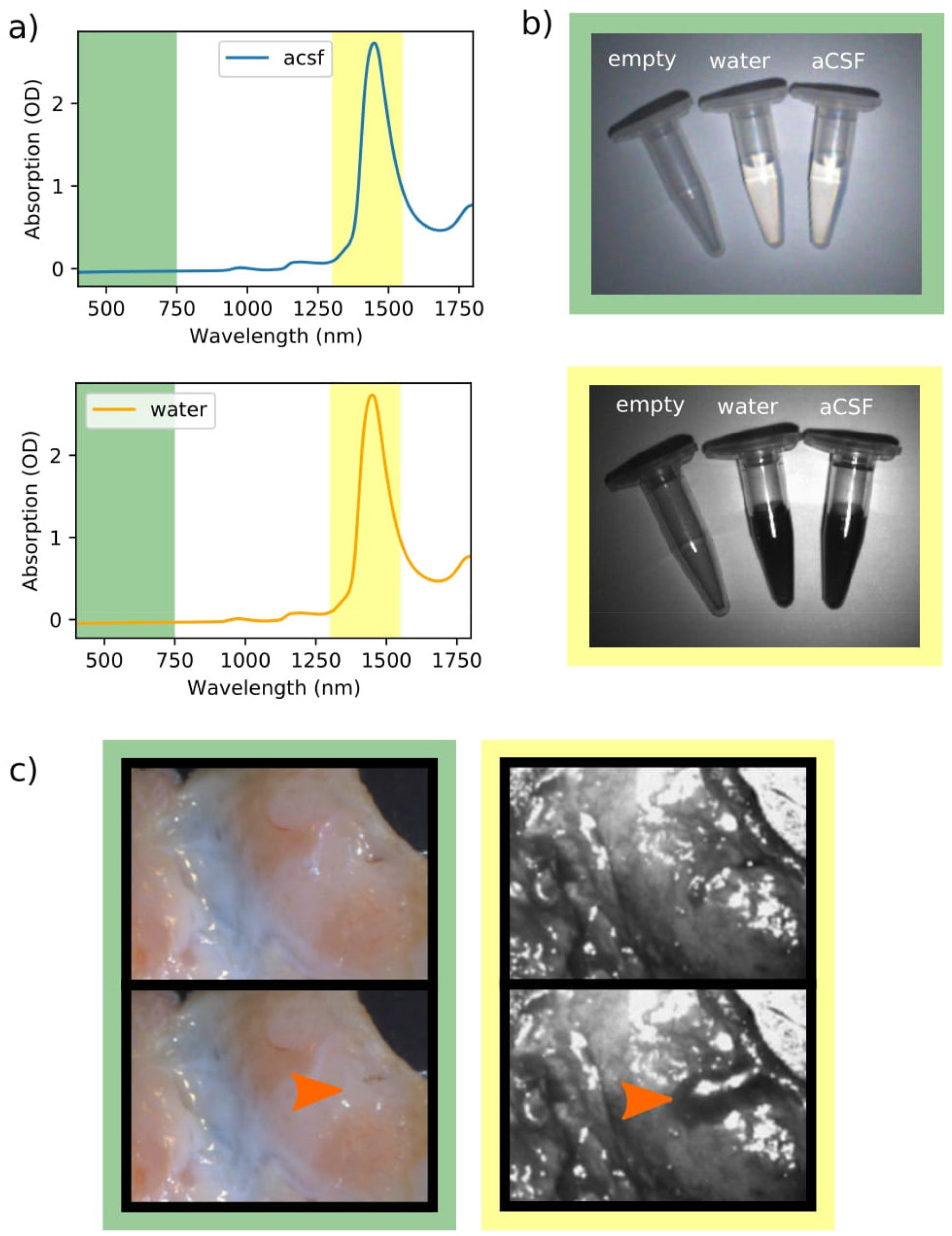
Optical properties of water and aCSF. a) Between 400nm and 1800nm there are no significant differences in the spectra of water and aCSF. b) Eppendorf tubes from left to right: empty, water, aCSF. Both water and aCSF show strong absorption at ∼1450nm, which causes them to appear black whereas they are translucent in the visible wavelength range. c) Comparison of tissue without and with the presence in water. Whereas in the visible spectrum the presence of water cannot clearly be defined, when using the broadband 1450nm SWIR imaging the water produces a much clear contrast to the surrounding tissue and is clearly visible as a dark area. The orange arrows indicate the area where water is present. The background colors correspond to the highlighted wavelength ranges in the spectra.

**Fig. 3.**
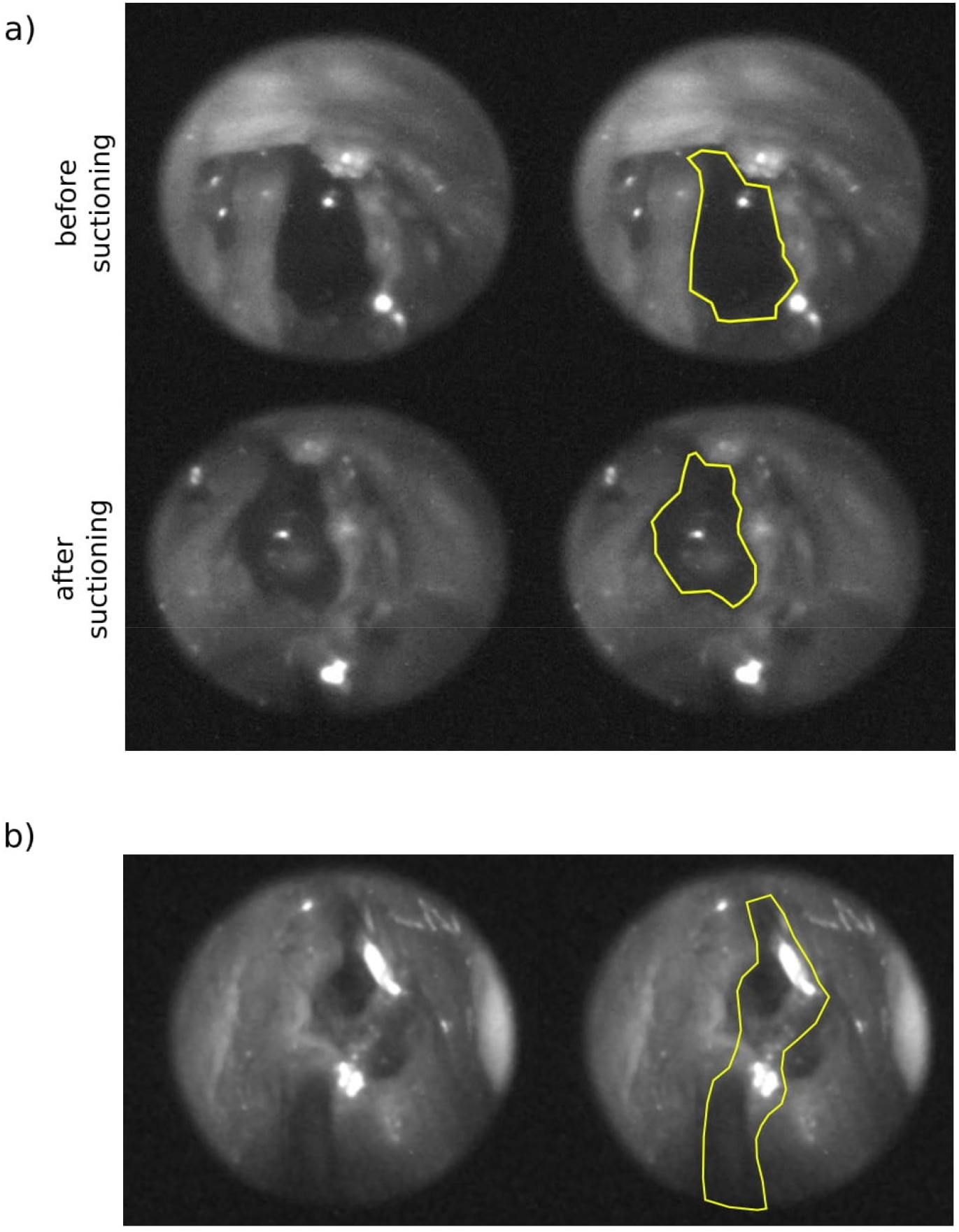
Imaging the skull base of a cadaver using the SWIR endoscope. a) The endoscopic SWIR images show that larger accumulations of water can be detected due to the increased amount of absorption. The yellow outline shows the cavity in which water accumulated, the left and right SWIR images are otherwise identical. b) Detection of thin films of water is difficult as there is not enough contrast between the thin films and the surrounding and underlying tissue. In the right image we have highlighted the area where water is present. For comparison, the left image is identical to the right, other than the water not being outlined. This shows that accurate detection of thin films of water was not possible with the endoscope design used for the cadaver study, which incorporated a 1450nm LED without an excitation filter.

**Fig. 4.**
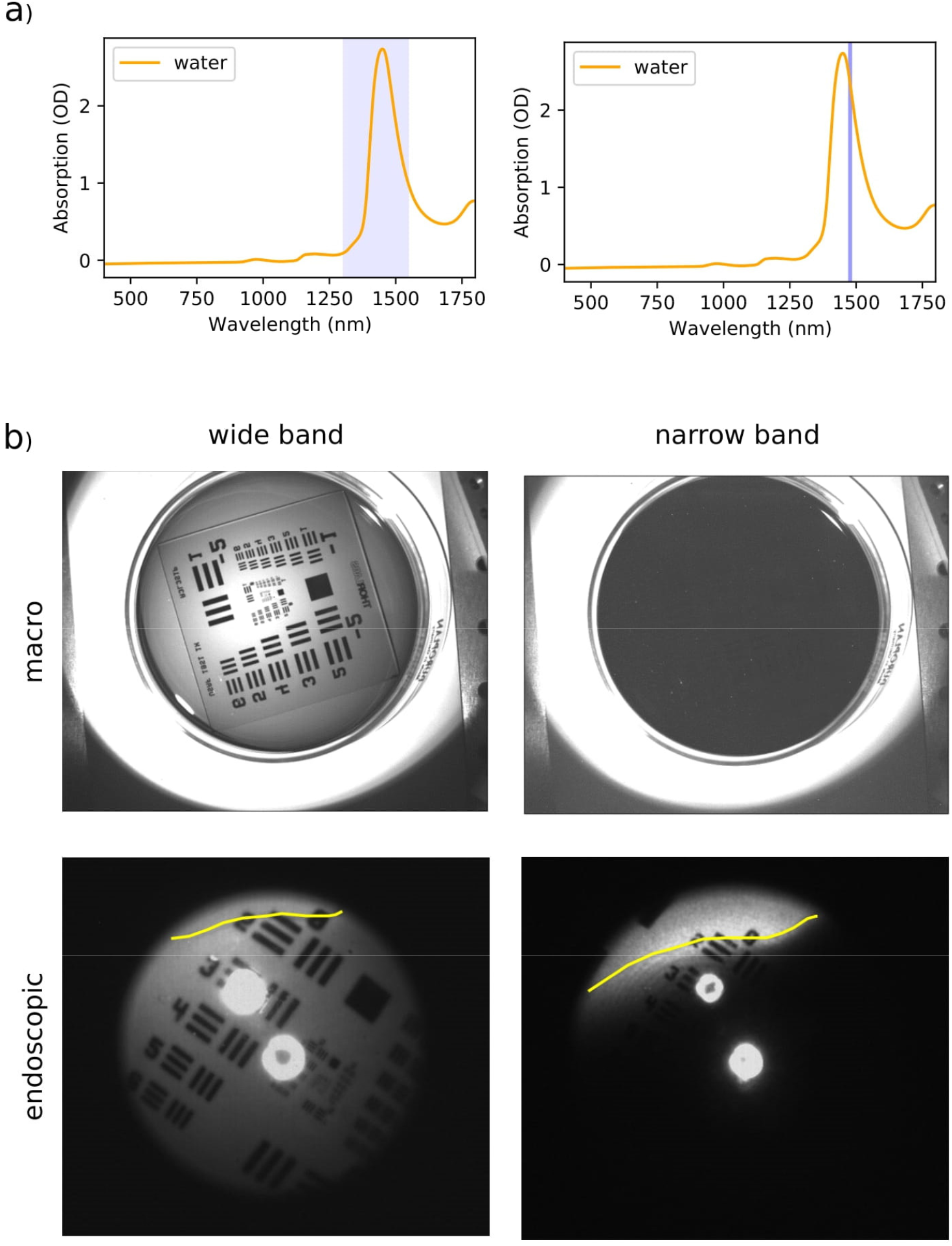
Showing the effect of a narrower wavelength range on the absorption (and thus detection) of water. a) The colored bands in the spectra show the wavelength range for the illumination of the images. b) When narrowing the wavelength range the absorption of the water clearly increases, making the water appear darker and less translucent. This becomes very apparent when viewing the test target with the different wavelength ranges, and the effect is present both in a macro imaging setup as well as in the endoscopic imaging setup. The yellow line on the endoscopic images shows the edge of the water.

**Fig. 5.**
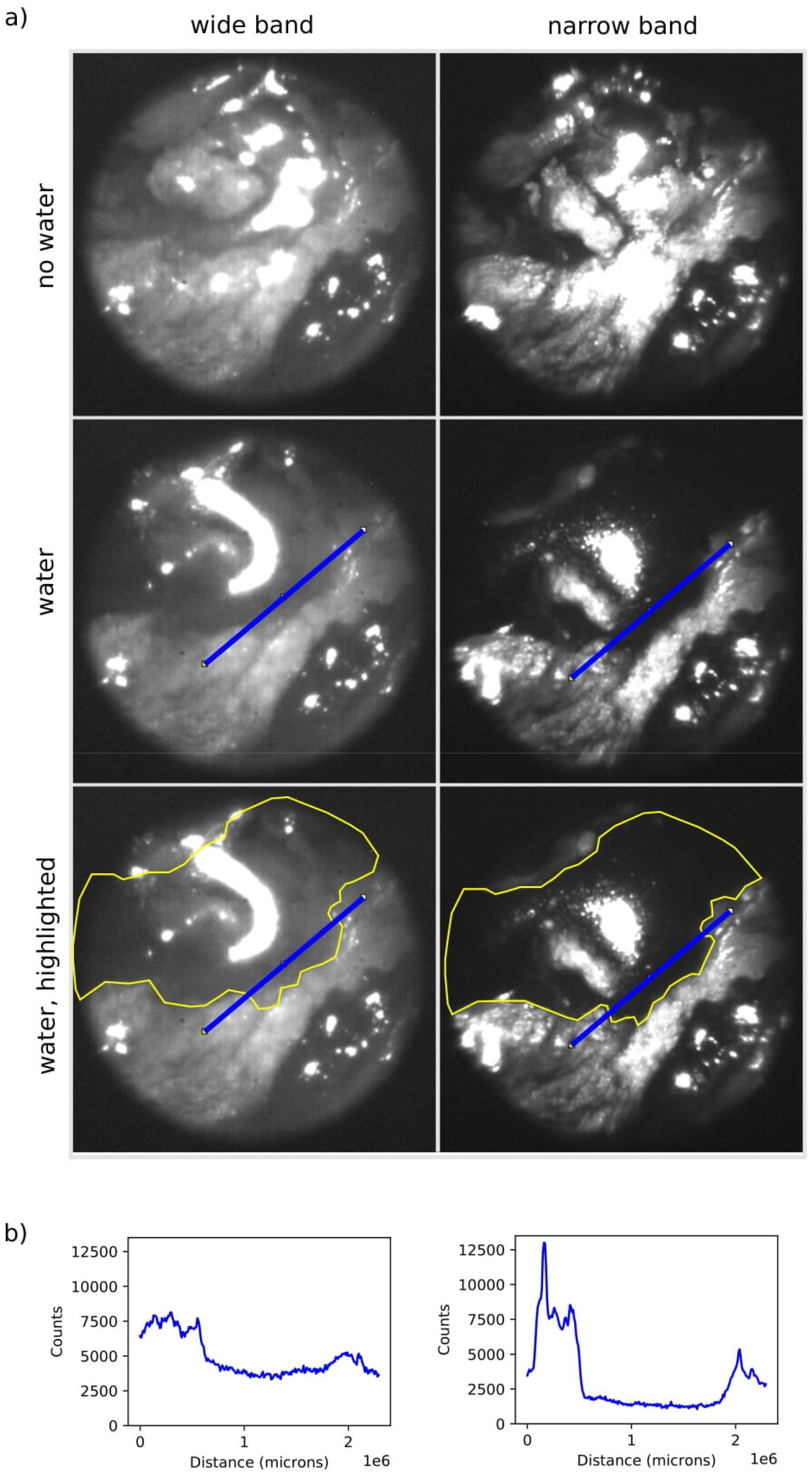
Showing the effect of a narrower wavelength range when imaging tissue. a) The narrowing of the wavelength range increases the overall contrast of tissue structures. It also makes it easier to pick out water on top of tissue, and it has an especially large effect on the edges and thinner areas of water. These images were taken without the use of polarizers. b) the graphs show the plot of the blue line profiles seen on the images above. The line profile cuts the edge of the water area. In line profiles over the water, we see that the narrow band image has higher contrast than the broadband image.

**Fig. 6.**
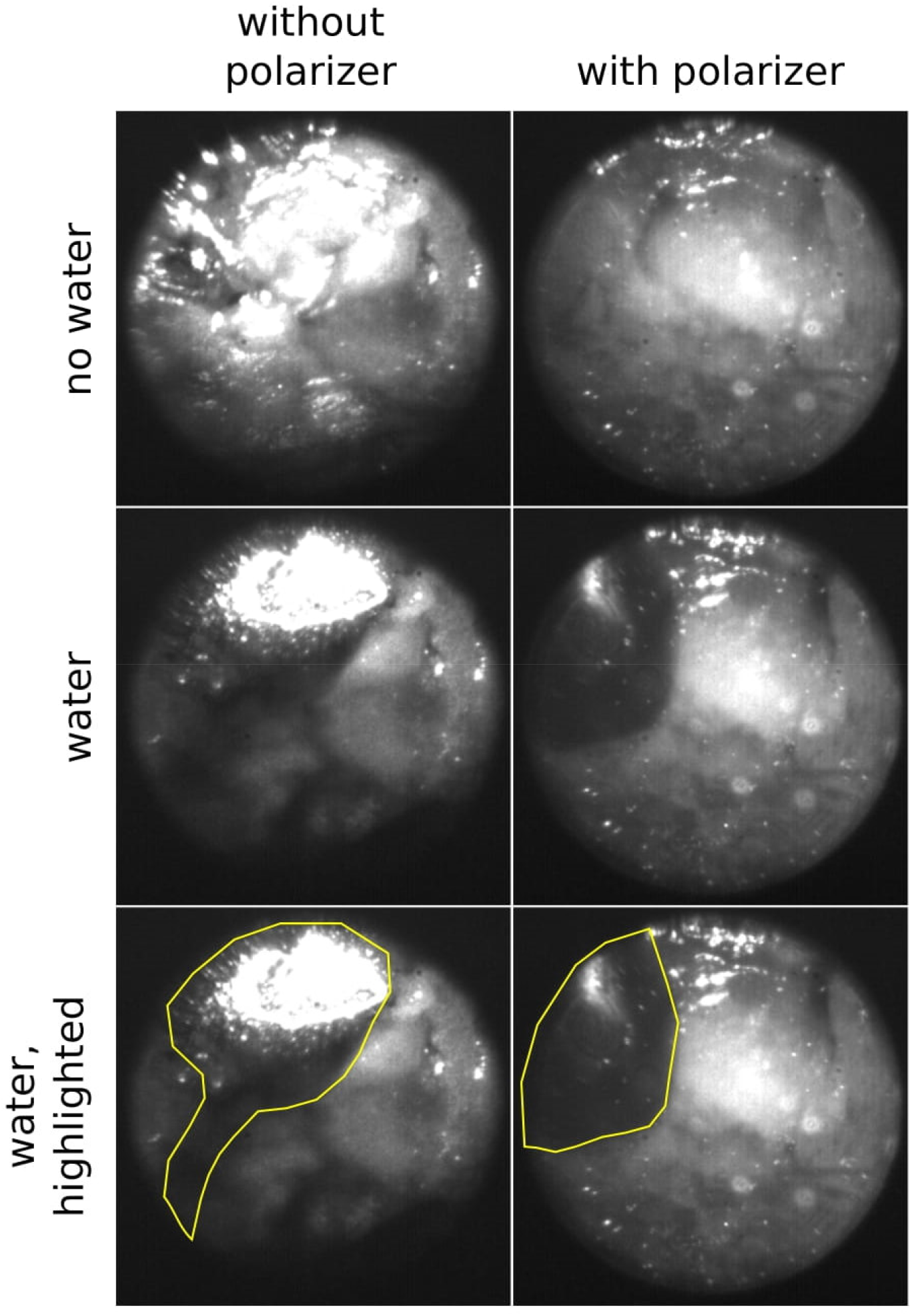
Comparison of imaging with and without the use of polarizers. The use of polarizers significantly reduces the overexposed areas in the image. This means that less of the areas of the image need to be omitted when analyzing and the presence of water in the image can be labeled with more certainty. The images were taken using the narrow band 1480nm laser for illumination.

**Fig. 7.**
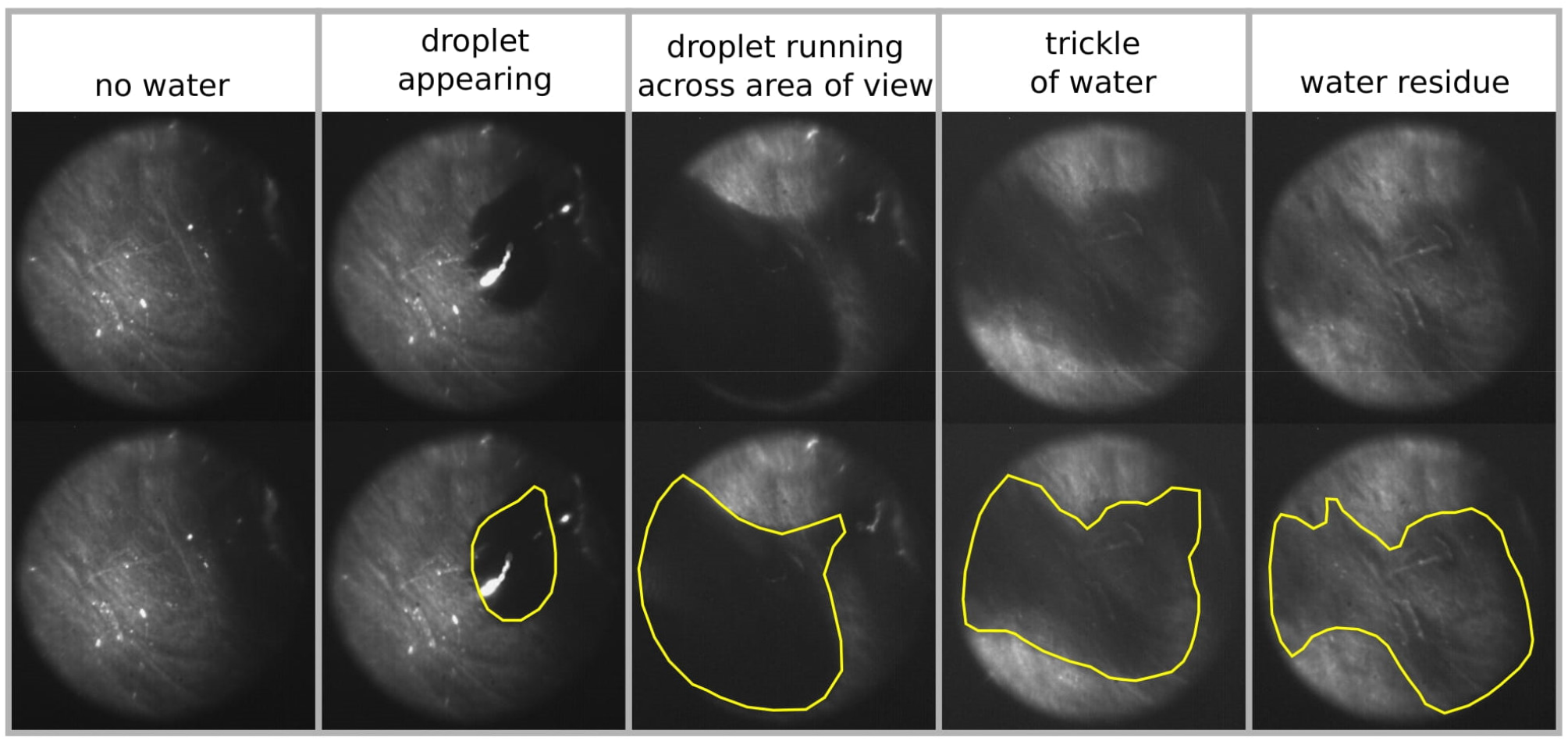
Images of the nasal cavity showing: no water, droplet appearing, droplet running across area of view, a trickle of water following in the path that the droplet left, residue of water. The bottom image shows the water highlighted and is otherwise identical to the image above it. With the revised endoscope design (including the polarizers and the narrow wavelength illumination) we significantly improved upon the quality of the data compared to our initial cadaver imaging. There are now fewer distracting reflections present, enabling us to clearly analyze almost the entire image and needing to omit very little areas in the image from analysis. Not only large droplets but also thin films and even small residues of water can be picked up against the tissue. An issue that needs to be addressed in further work is differentiating between shadows and water. In all images it is not possible to accurately diagnose water in the areas that are not well illuminated.

The imaging of the Eppendorf tubes confirmed that while being transparent in the visible there was strong attenuation when moving the 1450nm SWIR imaging window, for both water and aCSF. Whereas in the visible the fluid in the Eppendorf tubes is translucent, it appears dark and opaque in the SWIR and has a stark contrast to the air-filled Eppendorf tube and the background. Having confirmed that the optical properties of water and aCSF in the SWIR are near-identical we will use water for all experiments from this point on, due to water being readily available.

Since skin and other biological tissues have a high water conten,t we were expecting a reduction in contrast between water and background when moving to imaging the water on top of tissue. We tested whether moving to the SWIR would give us an advantage in the detection of water on top of tissue compared to white light imaging by using a piece of chicken skin (food grade; inner side of skin facing upwards/ towards camera). Using a piece of plastic tubing inserted through the chicken skin water could be pooled onto the surface of the chicken skin. The resulting images showed that in the visible there is no significant difference to be seen between chicken skin with and without water. In the SWIR however, the water droplet on top of the chicken skin can clearly be depicted against the surrounding skin.

### 3.2 Cadaver Study

The cadaver imaging confirmed that we can detect an accumulation of artificial CSF inside the human nasal cavity using the SWIR endoscope. The SWIR endoscope system is also able to show anatomical landmarks comparable to scopes currently used in sinus surgery. Detecting thin films of artificial CSF proved difficult, highlighting the need to make modifications to the system to be able to detect smaller amounts of liquid.

### 3.3 Use of a narrow-band SWIR laser improves CSF contrast

Previous data had shown that droplets of water and thick films of water gave good contrast to surrounding tissue, thin films on the other hand were much harder to depict as they showed too little absorption to contrast clearly against the background. We aimed to improve this by moving the illumination window to a narrower range and focusing it more on the attenuation peak we had seen in the spectroscopic data. The resulting images showed that narrowing the spectrum had a strong effect on the transparency of the water. While it was clearly possible to depict the test target under the thin layer of water when using the broadband LED this was no longer possible when adding the bandpass filter.

We adapted these findings to our endoscopic setup and exchanged the previously used broadband LED (M1450L4, Thorlabs) for a 1480nm narrow band laser (L1480G1, Thorlabs). The laser allows for improved optical efficiency and provides additional power at the narrow band of 1480nm which we would lose with a broadband LED and filter setup. We analyzed the effect this had on the endoscopic image in a similar experiment to the one described above for the macro setup. We again used a partially water-filled petri dish laid on top of a test target. To better be able to assess the effect the depth of the water had on its transparency we slanted the petri dish to create a gradient of water thickness. We added a very small amount of dish soap to the water to reduce the surface tension of the water and thus create a thinner film of water. The resulting images showed that the laser gave comparable images to those where the LED spectrum had been narrowed using a bandpass filter, and that the laser spectrum was sufficiently narrow for our purposes. While the test target was clearly visible throughout the entire image with the broadband illumination it was only clearly visible for very thin films (<<1mm) when using the narrow band illumination. With an increasing thickness of the film of water it was no longer possible to perceive the test target.

The question that remained was whether this would lead to an increase in contrast between water and tissue, or whether the absorption of the tissue increased at similar levels as that of the water. We used the same endoscope setup as had been previously used to image the petri dish with water, only swapping the petri dish out for a piece of chicken skin. We imaged the chicken skin without water and while running a droplet of water across it. The resulting images showed that narrowing the illumination helped to depict where water was present on the sample. This confirmed that narrowing the illumination spectrum around the absorption peak of water increases the chances of being able to pick up smaller amounts of water and to better be able to pick up the water against the surrounding tissue.

### 3.4 Reducing reflections through the use of polarizers

In previous data, we observe that some areas of the images are overexposed, due to the presence of direct reflections of the illumination source off the sample. We aimed to reduce these bright reflections by adding polarizers to our endoscope design. One polarizer was positioned inside the video coupler (between endoscope and camera) and a second polarizer was attached to the tip of the endoscope. The polarizer at the tip of the endoscope needed to be custom cut in order to only cover the fiber ring of the endoscope and not the lens. The polarizers were oriented in such a way to each other as to bring the reflections to a minimum. Using this modified endoscope setup, we again imaged a piece of chicken skin, first without water and then while running water across it. Comparing the images we obtained with and without polarizers we can show that adding polarizers to the design gives us two positive effects: Firstly, by reducing the bright reflections we are able to analyze more of the image accurately. Before, the regions displaying bright reflections could not be analyzed, as it was impossible to tell whether the tissue was causing the reflections, or whether the surface of the water was causing the reflections. Adding the polarizers reduced the size of overexposed areas on the images, thereby increasing the amount of information that could be obtained from one image. Secondly, reducing the bright reflections allowed us to increase the overall illumination, and this in turn enhanced the contrast between tissue and water.

### 3.5 Imaging the nasal cavity

For the final assessment of the improved SWIR endoscope design we used a ½ pig head (food grade) and imaged the nasal cavity. The endoscope setup was identical to the setup described in section 2.1. This setup combines the use of a narrow band illumination with the polarizers. Water was dripped into the nasal cavity near the skull base and trickled towards the tip of the nose. The nasal cavity was imaged without water present and while the water was trickling along it.

Using the pig head, we were able to show that we were able to detect both droplets of water and thin films of water inside a nasal cavity. Even water residue left behind on the tissue could be detected, showing that we were able to greatly improve upon our initial endoscope design.

## 4 Discussion

CSF leaks can occur in an iatrogenic, traumatic, or idiopathic setting. If not identified in a timely manner, they can lead to complications ranging from headaches to meningitis. Current options for CSF leak diagnosis include the off-label use of intrathecal fluorescein and subsequent visualization with nasal endoscopy, cisternogram with intrathecal contrast, or testing for the CSF specific compound Beta-2 transferrin (Lobo, Baumanis, & Nelson, 2017). Each method has its drawbacks. Both fluorescein and contrast require intrathecal delivery via a lumbar puncture (LP), which can cause significant discomfort and post-LP headache. Additionally, fluorescein injection has been associated with rare but serious side effects including paraparesis, numbness, and seizure (Raza et al., 2016). Moreover, the absence of fluorescein visualization can have a false-negative rate of as high as 26.2% in some studies (Seth, Rajasekaran, Benninger, & Batra, 2010). Beta-2 transferrin, on the other hand, requires ample volume of rhinorrhea to be present and often has a 24-72 hour delay for results (Haft, Mendoza, Weinstein, Nyunoya, & Smoker, 2004). Thus, a more accurate, immediate, and accessible means of intraoperative CSF leak detection is needed. We propose the use of a surgical system using an endoscope in the short-wave infrared region for endonasal assessment of CSF leaks. This technology offers the opportunity to diagnose CSF leaks quickly, at the point of care, and with minimal morbidity.

Prior to developing an endoscopic system, we obtained spectra of CSF, artificial CSF and water which were found to be identical. Because CSF is 99% water having a large absorption in the SWIR band around 1400-1500 nm, we are confident that this device has the potential to be an effective means of diagnosis. We have shown that it provides sufficient contrast between the CSF (which appears as black since it readily absorbs SWIR light) and the surrounding tissue. Based on the absorption spectra, we customized an endoscope with those wavelengths characteristics into a form factor used on endoscopic sinus surgery. It was tested on a 3D printed model simulating the nasal cavity and sinuses.

Testing our design on a cadaver showed that when using a broadband 1450nm LED light source we were not able to meet our criteria for CSF detection. The poor water absorption of thin layers of fluid showed diminished contrast and while illuminating the target area directly, there was a significant amount of reflections which reduced our ability to accurately detect artificial CSF. Modifications on our illumination source included incorporation of a laser with a narrow band wavelength of 1480 nm to enhance contrast from water absorption. To decrease the reflection artifact, we incorporated polarizers which improved our detection accuracy. This was corroborated using porcine nasal cavity mucosa.

To be able to use the endoscope in a clinical setting, further modification will be needed. The size and weight of the camera are important (and potentially limiting) factors, as both can interfere with the handling and ease of use of the endoscope. However, the endoscope is compatible with any standard c-mount SWIR camera, and small and lightweight SWIR cameras are increasingly becoming commercially available. Furthermore, while significantly improving image quality, the current polarizer setup has drawbacks on handling and ease of use of the endoscope. The polarizers would need to be revised and ideally manufactured directly into the endoscope in such a way that they stand up to the rigorous cleaning and disinfection needed for clinical use. Further, we would incorporate the polarizers into the video coupler in such a way that the camera can be freely rotated without impacting the polarizer orientation.

Limitations of our current study include the inability to image both on the visible and shortwave infrared which would be required for surgical procedures. Another limitation of our study is that our device has only been tested in cadavers and animal models which although valid testing models it does not represent the live scenario where blood and mucous secretions can be present. Moreover, currently used endoscopes and imaging systems for skull base surgery incorporate infrared imaging for ICG visualization to monitor perfusion of septal flaps.

Our future work will look to incorporate the ability to image both on the SWIR and the visible and also to incorporate fluorescence imaging while maintaining a weight and form factor that allows use in the operating room and can tolerate the rigors of surgical sterilization procedures.

## Acknowledgements

We also acknowledge funding from the Helmholtz Zentrum München, the National Center for Tumor Diseases Dresden (NCT/UCC Dresden), the Helmholtz Imaging Platform, the DFG— Emmy Noether program (no. BR 5355/2-1) and SFB1123, from the CZI Deep Tissue Imaging (DTI-0000000248) and BMBF (BetterView) to O.T.B. and funding from a Helmholtz Enterprise Grant to T.S.B. We thank M. Warmer and P. Tsrunchev for the excellent technical support. We also thank M. Warmer and A. Chmyrov for helpful discussion

## Author Biographies

**Tjadina Klein** is a PhD student at Helmholtz Pioneer Campus at Helmholtz Munich. She received her degree in veterinary medicine from the University of Leipzig in 2018.

**Tulio Valdez MD MSc** is an associate professor of Otolaryngology at Stanford University. He has an interest in photonic applications for diagnosis and treatment in the Head and Neck.

**Thomas Bischof PhD** is a senior scientist at the Helmholtz Pioneer Campus at Helmholtz Munich. He is focused on developing photonic methods for guidance and diagnostics.

**Oliver Bruns PhD** is a full professor at the Technical University Dresden and head of the department of Functional Imaging in Surgical Oncology at the National Center for Tumor Diseases Dresden (NCT/UCC Dresden). He is also a principal investigator at the Helmholtz Pioneer Campus at Helmholtz Munich.

**Michael Chang MD** is a clinical instructor in Rhinology and Skull base surgery at Stanford University.

**Stella Yang BS** is a research assistant at Valdez Lab.

**Mahbuba Tusty BS** is a medical student at Stanford University.

**Jayakar Nayak MD PhD** is a physician scientist and skull base surgeon at Stanford University.

